# Cobalt effects on prokaryotic communities living in growing river biofilms: impact on their colonization kinetics, structure and functions

**DOI:** 10.1101/2024.05.02.592147

**Authors:** Sarah Gourgues, Marisol Goñi-Urriza, Mathieu Milhe-Poutingon, Patrick Baldoni-Andrey, Nicholas Bagger Gurieff, Clémentine Gelber, Séverine Le Faucheur

## Abstract

1.

Although cobalt (Co) is widely used in the transition to low-carbon energy technologies, its environmental impact remains almost unknown. This study examines Co impacts on the prokaryotic communities of river biofilms to assess their potential use as bioindicators of Co contamination. To that end, biofilms were grown on blank glass slides placed in artificial streams enriched with Co (0.1, 0.5 and 1 µM Co) for 28 days and prokaryotic abundance and diversity were analyzed using DNA-metabarcoding every 7 days. The resilience of the prokaryotic community was investigated after a further 35 days without Co contamination. Prokaryotic communities were impacted by 0.5 and 1 µM Co from the beginning of the biofilm colonization. Although biofilms reached similar biomasses regardless of Co concentration, control biofilms were dominated by *Cyanobacteria* and *Planctomycetes* while *Bacteroidetes* dominated Co contaminated biofilms. Potential functional redundancy was observed with the implementation of carbon fixation alternatives by non-photosynthetic prokaryotes in biofilms subjected to high Co concentrations. No structural resilience of the biofilms was observed after 35 days without Co contamination. The use of prokaryotic community response measured using molecular approaches appears to be a promising and cost-effective approach for assessing changes in water quality due to metals.

**Graphical abstract:** 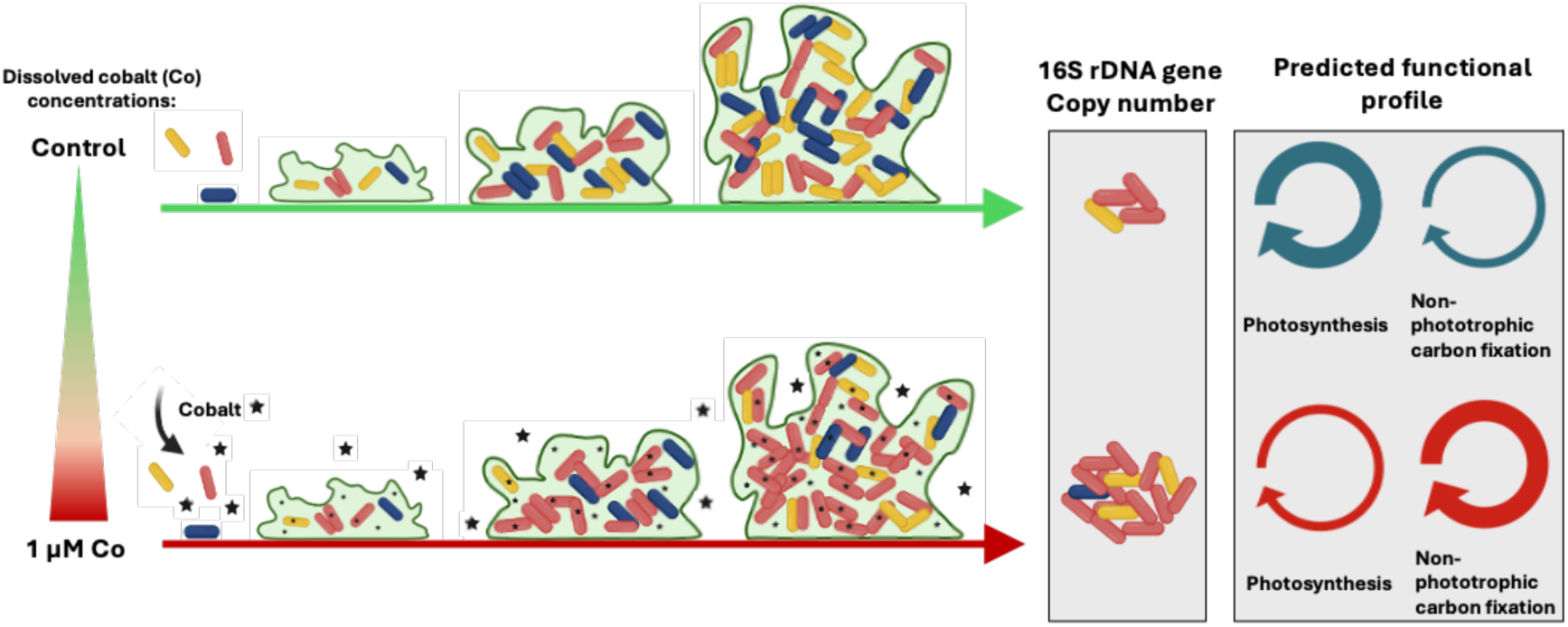

**Synopsis:** Few knowledge is available about Co ecotoxicity in freshwaters. This study assess the potential of prokaryotic communities developing in freshwater biofilms to be used as bioindicator of Co contamination.

## 2 Introduction

In the context of the energy transition, cobalt (Co) has become a metal of interest for the production of batteries. The increasing demand for lithium-ions batteries for vehicles and electronical devices led to an exceptional increase of its extraction (+95%) and its use (multiplied by 4) between 2008 and 2018^1–3^. Consequently, Co is nowadays regularly found with concerning concentrations in aquatic systems, especially near industrial sites^4,5^. As such, if background dissolved Co concentrations in European freshwater environments range from 0.17 x 10^-3^ to 0.27 µM Co (mean 5.65 x 10^-3^ ± 0.02 µM Co, n = 807)^6^, concentrations as high as 53.68 µM and 48.05 µM Co can be measured in mining effluents in Democratic Republic of Congo and near a Spanish industrial plant’s discharge, respectively^4–7^. However, environmental quality standards are still rarely available, such as in Europe or the United States. In Canada, the site-specific Federal Water Quality Guideline ranges from 0.01 to 0.03 µM Co depending on water hardness^8^. Its *Predicted No Effect Concentration* (PNEC) has been determined to be 0.02 µM (1.06 µg.L^-1^) in freshwaters^9^.

Cobalt is an essential micronutrient for organisms involved in numerous enzymatic processes^10^. However, studies performed on model organisms have demonstrated its toxicity at high concentrations. For example, the model bacteria *Geobacter sulfurreducens* had higher generation times when grown at Co concentrations equal or above 250 µM^11^. Decrease in pigment contents and growth rates have also been reported in microalgae in the presence of Co at concentrations equal to or above 170 µM^12,13^.

Aquatic biofilms are ubiquitous assemblages of microorganisms, which include bacteria, archaea, microalgae, fungi and meiofauna embedded in an extracellular polymeric substance (EPS) matrix^14–18^. This aggregate form of microbial life develops on submerged substrates and composes the basis of the trophic chain in aquatic ecosystems, especially in lotic systems. Their sensitivity to water quality changes makes them early warning tools of water quality degradation. Indeed, several studies have tackled the long-term consequences of metal contamination on biofilms, even after a recovery period without metal stress^19,20^. Up to now, few studies have focused on Co effects on biofilms. A study demonstrated that Co bioaccumulation was associated with a disruption of chlorophyll *a* synthesis while EPS production was enhanced^21^. Also, it was demonstrated that prokaryotic and microeukaryotic communities’ compositions and meta-metabolome from mature biofilms translocated in Co contaminated waters were rapidly impacted in comparison with natural biofilms^22^. These studies, led mature biofilms have not dealt with Co impact on biofilm colonization and setup yet. However, high vulnerability of biofilms at early stages of colonization were demonstrated in the presence of Cd^23^ or TiO_2_ nanoparticles^24^.

Taxonomy and abundance analyses of diatoms living in biofilms is currently part of the European water quality assessment, while not takin into account the prokaryotic community of the biofilm. Indeed, while rarely used in water quality assessment, prokaryotes are known to respond quickly to changing environmental conditions, including change in metal concentrations. They also provide numerous ecosystem services in collaboration with other organisms. For example, auto- and heterotrophic organisms are strongly linked, and their metabolic cooperation is essential for major element cycling^25^. That dependence between communities defines biofilms as a functional consortium, on which stresses will have repercussions on the overall microecosystem structure and functioning. The improvement of molecular techniques in the last decades, with the permanent amelioration of 16S rDNA gene databases has facilitated a more holistic comprehension of prokaryotic response to stresses, encompassing both the community structure and functional profile. Hence, prokaryotes may represent a more rapid and complete alternative to microalgae analysis for bioindication of metal contamination in freshwaters.

The aim of this study was to examine the effects of Co on prokaryotes in a growing biofilm to assess their possible use as bioindicators of metal exposure in aquatic ecosystems. To do this, biofilms were grown in artificial streams contaminated with Co and analyzed for its prokaryotic community abundance, structure, and functional potential after 7, 14, 21 and 28 days of exposure. The prokaryotic community was also investigated after a recovery period (without contamination) to investigate the resilience of the biofilms.

## 3 Material and methods

### Exposure experiments and sample collection

Exposure experiments were undertaken in outdoor artificial streams at the Pilots Rivers facilities (TotalEnergies, Lacq, France). The artificial streams were continuously filled by the *Gave de Pau* river without any pre-treatment at a flow rate of 7.5 m^3^.h^-1^ (Figure S1). Cobalt (Cobalt(II) chloride hexahydrate, 98%, Thermo Scientific Chemicals) was added to nine streams to reach nominal concentrations of 0.1 µM, 0.5 µM and 1 µM in triplicate (3 treatments × 3 streams). These Co concentrations were 5 to 55 times higher than the PNEC (0.02 µM)^9^, representing thus environmentally relevant concentrations. Three additional streams were used as control environments, which were only filled with the Gave de Pau water without Co addition.

At the beginning of the exposure experiments (Day 0), thirteen sterile blank glass slides (10 x 20 cm) were placed in each stream to serve as the support for biofilm colonization. Water and biofilm were collected every seven days during biofilm colonization (D7, D14, D21 and D28) as well as 35 days after stopping the addition of Co, the so-called the Recovery period (DR). Slide triplicates in each channel were scratched at each sampling event during Co exposure. Only one slide was used after the DR.

On the day of sample collection, the main physico-chemical parameters of the water, i.e, pH, temperature, dissolved oxygen, and conductivity, were measured in each stream. Water was also collected and stored at 4°C for the measurements of alkalinity, dissolved cations, including Co, anions and organic carbon concentrations (Text S1). Cobalt speciation was calculated using the Windermere Humic Aqueous Model (*WHAM,* version 7.05.5)^26,27^. Biofilms were collected in 2 mL sterile Eppendorfs by scraping the colonized glass slides with sterile microscope slides. Detailed illustrations of glass slides collected for biofilm sampling are presented in Figure S2. Samples were weighed and stored at -80°C for molecular analyses and at -17°C for ancillary parameters analyses. Biofilms dry weight and chlorophyll content were measured for each sample and more detailed protocols are detailed in (Text S2).

### Cobalt bioaccumulation

Total and intracellularly bioaccumulated Co were measured in freeze-dried biofilms mineralized with a mixture of nitric acid 70% (metalgrade) and hydrogen peroxide (H_2_O_2_; ultratrace) 30 % (v/v; 2:1) in a UltraWAVE oven (Milestone, Italy). For intracellular analysis, fresh biofilms were firstly rinsed with 10 mM of ethylenediaminetetraacetic acid (EDTA >= 99%, Sigma-Aldrich, Germany) for 10 min and rinsed with filtered (0.45 µM) water from the control stream of pilot rivers^22,28^. Cobalt concentrations in the mineralized biofilms were measured using ICP-MS (Agilent ICP-MS 7500, CA, USA). The certified standard reference material (BCR-414) (Plankton, JRC, Brussels) was used to verify the biofilm digestion (recovery of 86.3 ± 8.6 %, n=6).

### Diversity of prokaryotic communities

DNA extractions were performed using PowerSoil DNA Isolation Kit (Qiagen, Germany) following the manufacturer’s instructions. Extracted DNA was quantified using a Qubit 1X dsDNA BR assay kit (ThermoFisher Scientific, Country). The V4-V5 hypervariable regions of 16S rDNA gene were sequenced after an amplification with the universal primers V4-515F and V5- 928R (Wang and Qian, 2009). The PCRs were performed with a final reaction volume of 50 µL using 25 µL of ampliTaq Gold 360 master mix (Applied Biosystems, CA, USA), 0.4 µM of each primer and 1 µL of DNA template (between 7.2 and 412 ng). PCR amplification was carried out as follows: initial denaturation of 10 minutes at 95°C, 30 cycles of 30 s denaturation at 95°C, 30 s annealing at 65°C and 40 s of elongation at 72°C. Then a final elongation of 7 minutes at 72°C was performed. Amplicons were then sequenced by *La Plateforme Génome Transcriptome de Bordeaux (PGTB, Bordeaux, France)* using Illumina Miseq 2 x 300 pb method and reagent kit V3. Raw sequences were submitted to the National Center for Biotechnology Information Sequence Read Archive under the accession number PRJNA1098091.

Raw sequences were then processed using the FROGS (version 4.1.0) pipeline (Find Rapidly OTUs with Galaxy Solution)^29^. The merging, denoising and dereplication of the raw sequences were performed using VSEARCH tool^30^. Sequences with less than 200 bp were removed for further analysis. Clustering in operational taxonomic units (OTUs) was done with Swarm^31^, setting the aggregation distance clustering to 1. Chimeras and OTUs representing less than 0.005% of all sequences were removed^32^. Data was normalized to 10,675 reads per sample and the final total number of OTUs was 1437. Taxonomic assignments of OTUs were performed using 16S SILVA 138.1 and OTUs associated with chloroplast were removed for further analysis. Functional profiles of communities based on 16S sequences were predicted using PICRUSt2^33^. Resulting metabolic profiles were then annotated using Kyoto Encyclopedia of Genes and Genomes (KEGG) pathways with ggpicrust2^34^ package in Rstudio.

### Quantification of prokaryotes abundances

16S rDNA genes copy number was quantified using the DyNAmo Flash SYBR Green qPCR Kit (ThermoFisher Scientific, USA) with the same primer pairs as those used for microbial diversity (see above), without MiSeq adaptors. Quantitative PCR was performed in a final volume of 20 µL using 0.5 µM of each primer and DNA template (between 1.44 and 41.2 ng). Amplification was done as follows: after an initial denaturation of 7 minutes at 95°C, 40 cycles of 10 s of denaturation at 95°C, 30 s of annealing at 57°C and 30 s of elongation at 72°C were performed. A melt curve was produced to check the amplification specificity. Absolute quantification was performed with home-made standards consisting in pGEM cloned amplicons of *Escherichia coli* 16S rDNA gene. Analyses were performed in triplicates for each biological replicate. PCR efficacity was of 95.6%.

### Statistical analysis

Alpha diversity metrics of specific richness, diversity (Shannon index) and evenness (Pielou index) were calculated with vegan package on RStudio. All comparisons were assessed on Rstudio using Anova and Tukey post-hoc tests when possible. If conditions of application were not respected, a non-parametric approach was used with Kruskal-Wallis followed by Dunn post-hoc tests. Beta-diversity was evaluated with non-metric multidimensional (NMDS) ordination based on Bray-Curtis dissimilarity distances between OTU. Significant differences between communities were then assessed by permutational multivariate analysis of variance (PERMANOVA) with vegan package followed by pairwise comparisons with pairwiseadonis2 package. Comparisons between proportions of taxonomic groups (representing more than 1% of community) and functional pathways from PICRUSt2 analysis were performed using Statistical Analysis of Metagenomic Profiles (STAMP)^35^ applying White’s non-parametric test and an effect size filter for proportion differences >1%.

## 4 Results

### Physico-chemical parameters of waters

The physico-chemical parameters in the water streams were stable over the experiment except water temperature, which decreased from 14.30 ± 0.12 °C (Day 7) to 8.17 ± 0.03 °C (Day63) (Table S1). The average dissolved Co concentration in the control stream was 4.62 ± 4.02 nM. The average Co concentrations over the 28 days of exposure were consistent with the nominal concentrations (0.092 ± 0.004, 0.525 ± 0.084 and 1.089 ± 0.07 µM, respectively) (Table S1). Cobalt was predicted to be mainly complexed by carbonates (CoCO_3_) and under its free form (Co^2+^), representing 53 ± 3 % and 29 ± 2% of the total dissolved Co, respectively; the proportion of Co bound to fluvic acids being only 0.83 ± 0.51 % (Table S1). After the recovery period, dissolved Co concentration returned to its natural concentration in all of the streams.

### Co accumulation

Both total and internalized bioaccumulated Co were positively correlated with the free Co^2+^ concentrations (Figure 1 and Table S2), with Spearman’s Correlation coefficients between 0.82 and 0.97 (pvalue < 0.005). Throughout the experiment, total bioaccumulated Co in control and exposed biofilms remained stable, regardless of Co exposure concentration (Figure S3-A; Table S2). In contrast, internalized Co increased over time in Co-exposed biofilms (*p* < 0.05) (Figure S3-B), reaching 3.47 ± 1.63 µmol Co.g^-1^ DW for biofilms exposed to 0.1 µM of Co, 4.23 ± 0.25 µmol Co.g^-1^ DW for biofilms exposed to 0.5 µM and 7.68 ± 1.10 µmol Co.g^-1^ DW for biofilms exposed to 1 µM at D28. After the recovery period, bioaccumulated Co decreased in exposed biofilms (*p* < 0.05), but levels of Co remained between 4.35-fold and 8.68-fold higher than those found in the control biofilms (*p* < 0.05) (Figure S3-A,C).

**Figure 1:**
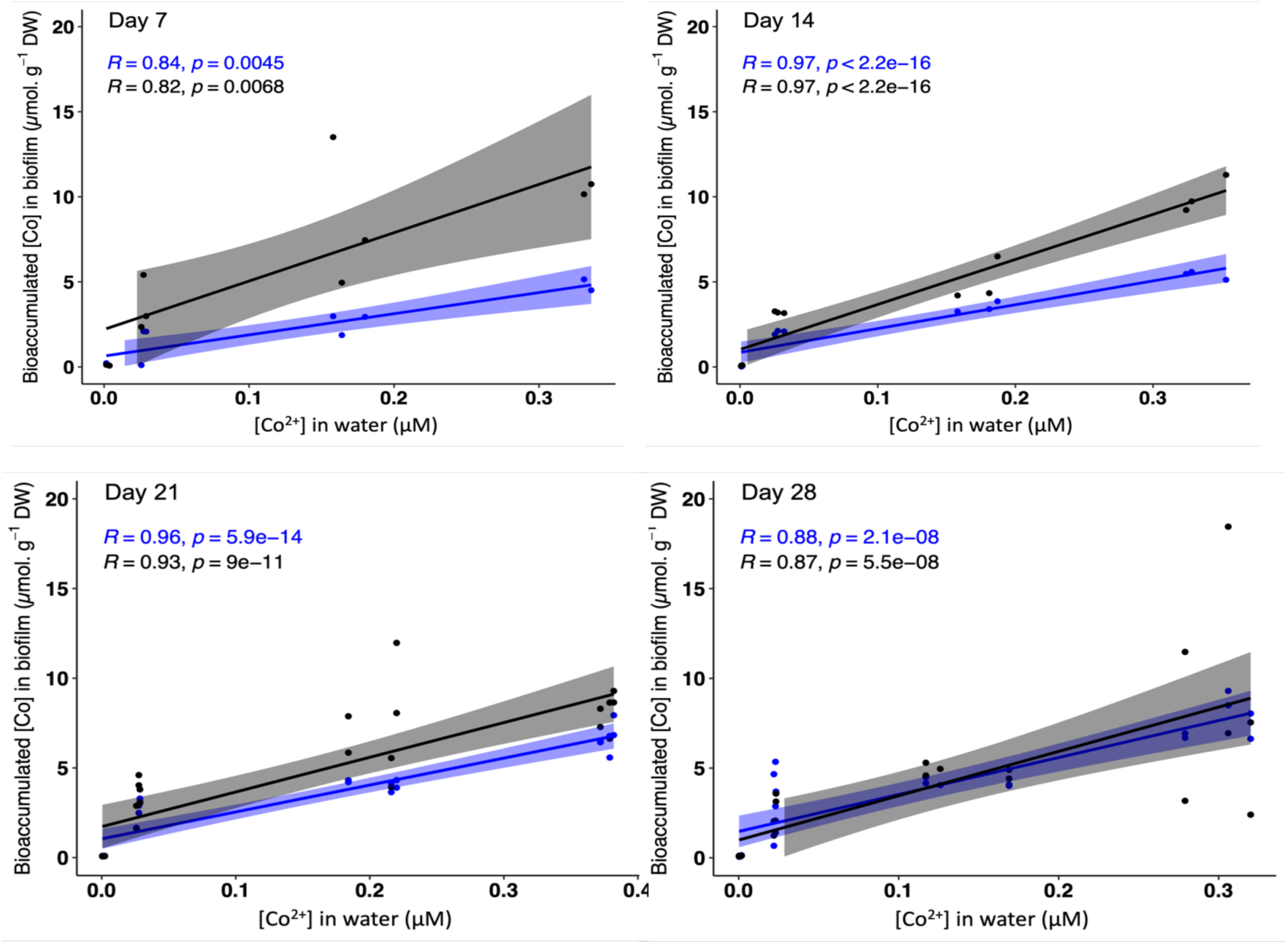
Cobalt bioaccumulation by biofilms over the colonization period. Lines represent Spearman’s correlation of total bioaccumulated Co (**black**) and internalized Co (**blue**) as a function of free ion Co^2+^ concentration in exposure medium at each sampling time.

### Biofilm structure as a function of Co exposure

Dry biomass of biofilms increased over colonization time with significant increases between D7 and D21, then a stabilization after that day, regardless of the Co exposure condition (Figure 2A, Table S3). Increasing levels of chlorophyll *a*, *b* and *c1 + c2* were measured in control and exposed biofilms (Figure 2B and Table S3). Higher content of chlorophyll *b* was observed in biofilms exposed to 1 µM of Co at D21 and D28 by comparison with control or 0.1 µM-exposed biofilms (*p* < 0.05). On the contrary, chlorophyll *a* and *c1 + c2* contents decreased as a function of Co exposure concentrations and were significantly lower in biofilms exposed to 0.5 µM Co by comparison with the control (Figure 2B, Table S3).

**Figure 2:**
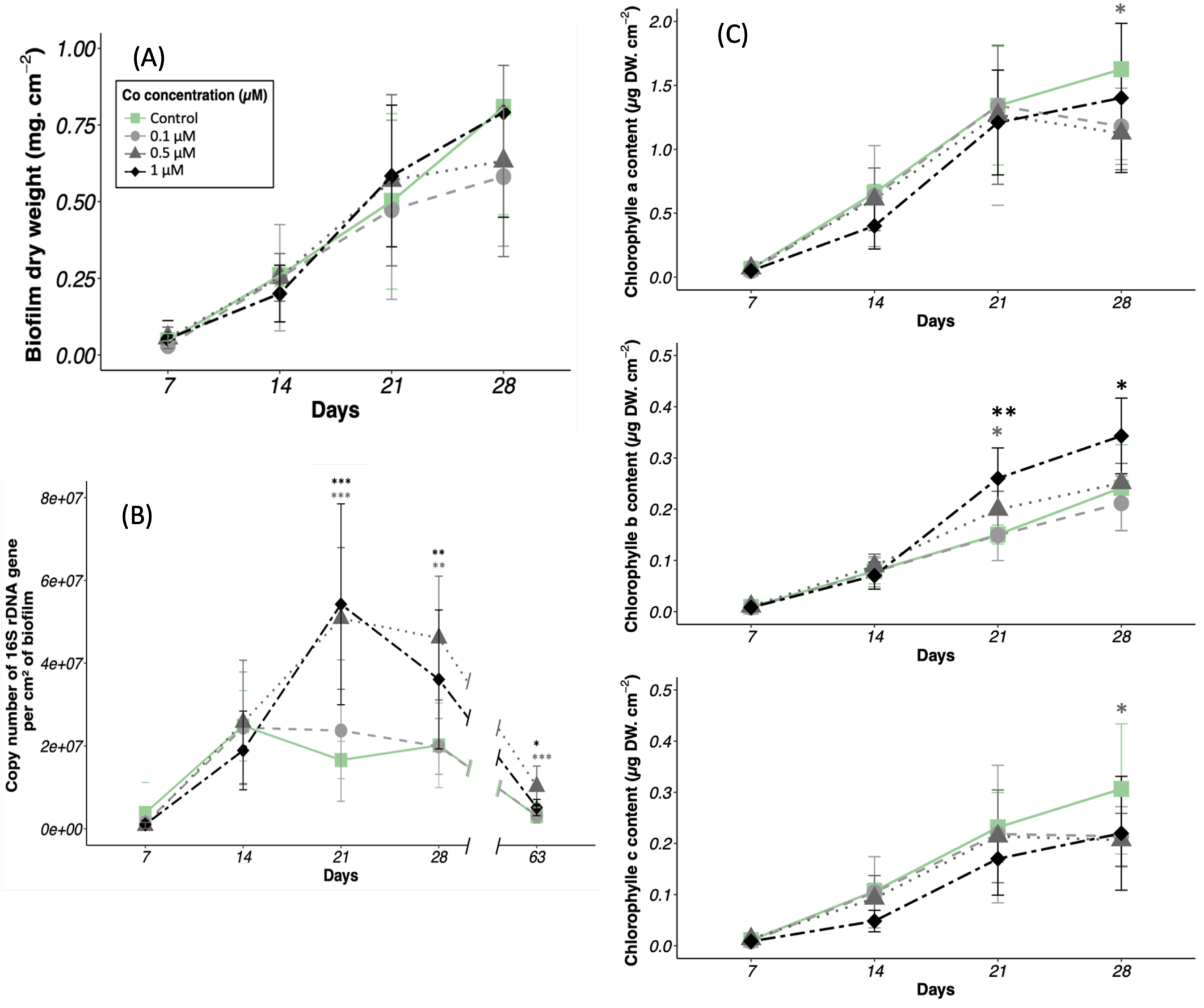
Main biological parameters of biofilm growing in the different artificial streams exposed to 0, 0.1, 0.5 or 1µM of Co over time. (A) Dry weight, (B) Chlorophyll *a, b and c* contents, (C) 16S rDNA gene copy numbers. Dry weight and chlorophyll content data includes samples in one replicate for Day 7 and Day 14, and two replicates within a stream for Day 21 and D28. For 16S rDNA gene copy numbers data includes samples in triplicates within a stream. * indicates statistical differences (*=p<0.05; **p<0.01), the color of lines indicates the increasing Co concentrations by comparison with control conditions (▪: Control conditions; • : 0.1 µM Co; ▲ : 0.5 µM Co; ♦: 1 µM Co)

### Prokaryotic abundance and diversity in biofilms

The number of 16S rRNA genes (used as a proxy of prokaryotic abundance) increased in all biofilms between D7 and D14 (*p* < 0.01), regardless of Co concentration (Figure 2C). After D14, the abundance of prokaryotes in control and 0.1 µM-exposed biofilms remained stable, but significantly increased when exposed to 0.5- and 1 µM of Co (*p* < 0.05). At D21, 0.5- and 1 µM- exposed biofilms had 3-fold higher levels of 16SrRNA gene copy numbers than the control or 0.1 µM-exposed biofilms (*p* < 0.05). 16SrRNA gene copy numbers significantly decreased between D28 and DR for all biofilms (*p* < 0.005). Nevertheless, after the recovery period, the prokaryotic abundances of biofilms pre-exposed to 0.5 µM and 1 µM of Co remained higher than the control biofilms (*p* < 0.01) (Figure 2C).

Control biofilms had stable specific richness figures during the 28 days of growing with 1004 ± 84 OTUs (Table S4). Nevertheless, both Shannon and Pielou indexes increased significantly at D28 (*p* < 0.05), indicating more evenly distributed communities (Table S4). On the contrary, specific richness of biofilms exposed to Co increased at D21, with significantly higher OTUs number and higher Pielou evenness for 0.5 µM exposed biofilms (*p* < 0.05) (Table S4). Specific richness, Shannon and Pielou indexes significantly decreased after the recovery period in most of the biofilms (*p* < 0.001) (Table S4).

The two-dimensional Non-Metric Multi-Dimensional Scale (NMDS) ordination was based on Bray-Curtis distances of prokaryotic communities clustered samples according to the growing time (axis NMDS 1) and Co concentrations (axis NMDS 2) (Figure 3). All biofilms, including the control, had changing communities over biofilm growth (*p* < 0.01) except biofilms exposed to 0.1 µM Co, which remained similar between D21 and D28 (Table S5). Control biofilms and biofilms exposed to 0.1 µM Co showed similar composition of the communities. On the contrary, biofilms exposed to 0.5 and 1 µM Co showed significant differences in the prokaryotic composition compared with control or 0.1 µM exposed biofilms, and also between them both, for all sampling periods (*p* < 0.01) (Table S5). Distances between these clusters increased with time, indicating that the dissimilarities between biofilms increased. At DR, the biofilms structures remained as different as they were during Co exposure without overlapping clusters (Figure 3) (*p* < 0.005) (Table S5).

**Figure 3:**
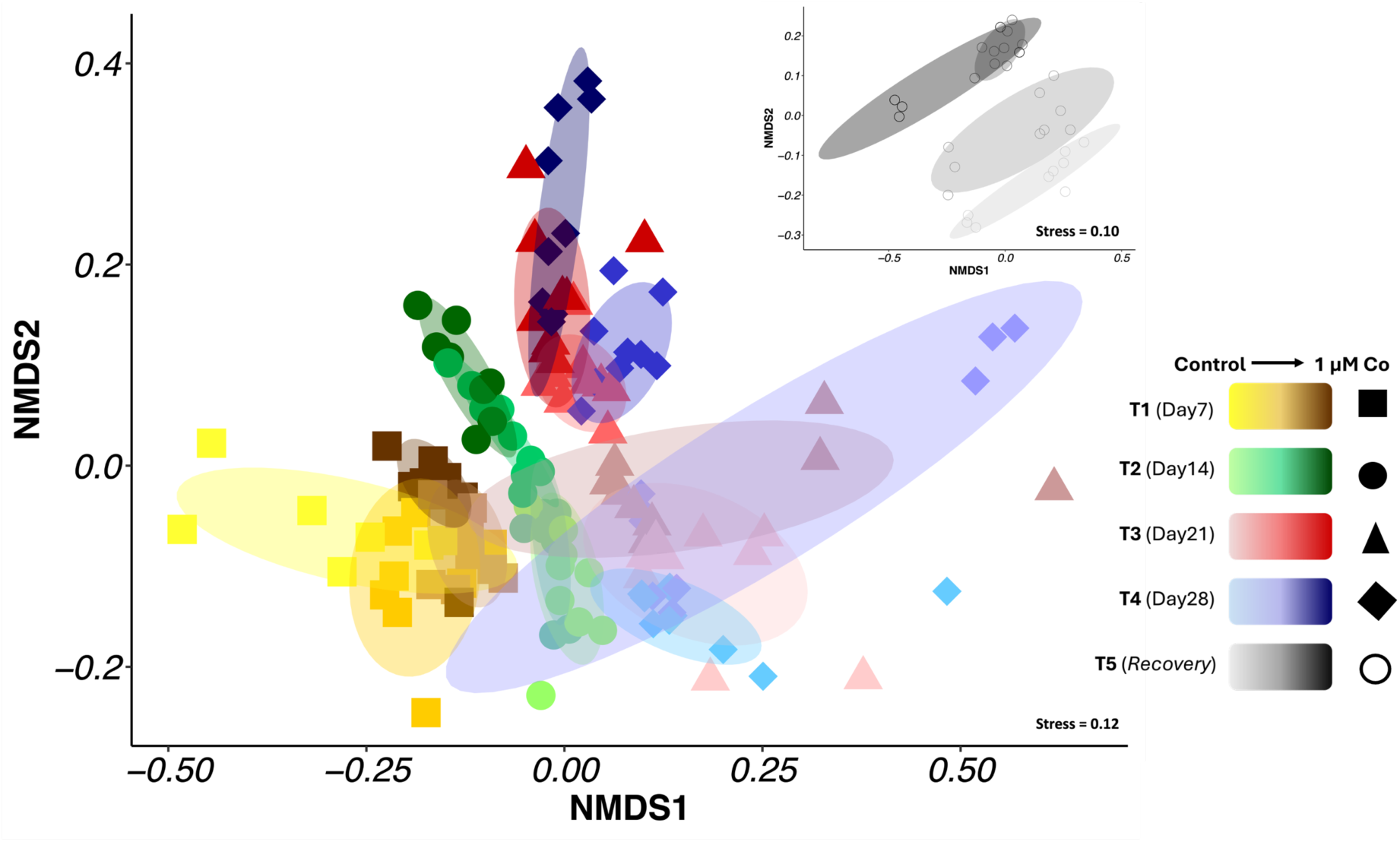
Non-metric Multi-Dimensional Scaling (NMDS) ordination based on Bray-Curtis dissimilarity matrix of OTUs from prokaryotic communities in growing biofilms in presence of Co (stress = 0.12). Insert corresponds to NMDS ordination of OTUs after *recovery* (stress = 0.10). Sampling time are represented as follows: ▪ T1 (Day 7); • T2 (Day14); ▲T3 (Day 21); ♦ T4 (Day 28) and **⭘** T5 (Recovery). Conditions of exposure are represented with a colour gradient from pale (control) to bright (1 µM Co).

Regardless of the time and Co exposure conditions, biofilms were dominated by *Proteobacteria, Bacteroidota, Cyanobacteria,* and *Planctomycetota* phyla (Figure S4). *Acidobacteria, Actinobacteria, Chlorofexi* and *Verrucomicrobiota* phyla were also found within biofilms but at a lower percentage. The prokaryotic community of control biofilms at D7 was dominated by *Cyanobacteria* (31.70 ± 25.00 %), with their relative abundance decreasing over time to reach 7.00 ± 4.04 % of the community at D28 (Figure 4B, Figure S4). *Bacteroidetes* followed the same trends, with a minimum observed at D21 (Figure 4B, Figure S4). On the contrary, the proportion of *Planctomycetes* class increased between D7 and D21 (*p* < 0.05) (Figure 4B).

**Figure 4:**
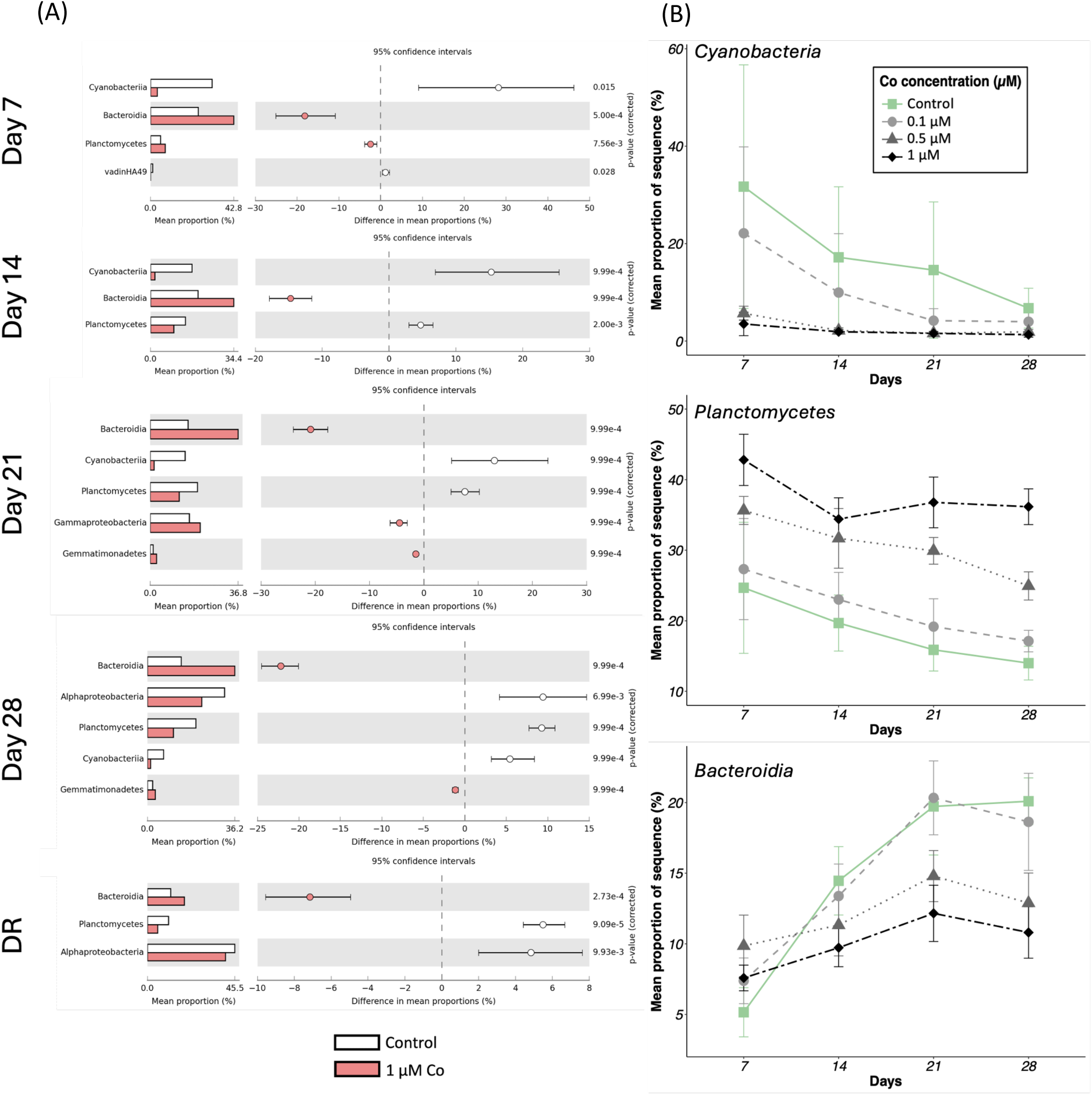
(A) Mean proportions and differences in mean proportions of taxa with statistically significant differences (White’s non-parametric test) between control and 1 µM-exposed biofilms based on STAMP comparison. In white, control condition, in red, biofilms exposed to 1µM of Co. (B) Modifications of relative proportions of *Cyanobacteria, Planctomycetes* and *Bacteroidetes* classes within biofilms. The color of lines indicates the increasing Co concentrations by comparison with control conditions (▪: Control conditions; • : 0.1 µM Co; ▲ : 0.5 µM Co; ♦ : 1 µM Co).

Cobalt exposure led to a modification of major prokaryotic taxa proportions in exposed biofilms (Figures 4A). As expected from the NMDS analysis, few modifications of the community composition were observed at 0.1 µM of Co (Figure S5), but the community significantly changed at 0.5 and 1 µM Co (Figure 4B, Figure S6). *Cyanobacteria* was highly sensitive to Co with significantly lower proportions in comparison with the control for all sampling points (*p* < 0.05). At D7, *Cyanobacteria* were already significantly impacted representing 5.7 ± 1.5 % and 3.5 ± 2.4 % of the communities for 0.5 µM and 1 µM-exposed biofilms, respectively (Figure 4B). *Planctomycetes* members were also negatively impacted at 0.5 µM and 1 µM Co from D14 to D28 (*p* < 0.05). On the contrary, *Bacteroidetes* was highly resistant, representing 42.8 ± 3.6 % and 36.2 ± 2.7% of the community at D7 and D28 at 1 µM Co. They represented only 24.7 ± 9.3% and 14.0 ± 2.5% at D7 and D28 in control biofilms (*p* < 0.05) (Figure 4B). The *Bacteroidetes* class was mainly composed of *Chitinophagales, Flavobacteriales* and *Sphingobacteriales* orders. At DR, biofilms which were exposed to 0.5 and 1 µM Co showed a significant increase in the relative abundance of *Cyanobacteria* (*p* < 0.001) and a decrease for *Bacteroidota* (*p* < 0.001).

### Functional profile of Co-contaminated growing biofilms

The predictive functional potential of biofilms was calculated on pathways related to biofilm formation and energy metabolisms (photosynthesis, carbon fixation pathways in prokaryotes, methane, nitrogen, and sulfur metabolisms). No variation on selected pathways was observed between control and biofilms exposed to 0.1 µM Co until D21, when photosynthetic and sulfur metabolism potential decreased whereas carbon fixation pathways inferred to prokaryotic communities were slightly enhanced (*p* < 0.05) (Figure S7). For higher Co concentrations (0.5 and 1 µM Co), pathways linked to carbon fixation were influenced by Co from an earlier stage of colonization (D7) (*p* < 0.05) (Figure S8, Figure 5). Indeed, significantly lower photosynthetic potential was observed but alternative prokaryotic pathways for carbon fixation were enhanced during the whole period of exposure (*p* < 0.05). Pathways linked to biofilm formation (including the attachment mechanisms and EPS secretion) were also negatively impacted in the presence of 0.5 and 1 µM Co from D7 (*p* < 0.05). Similar trends were observed for D14, D21 and D28 (Figure S8, Figure 5). Significant variations were observed for other energy metabolisms such as for methane, sulfur and nitrogen (*p* < 0.05) but without any clear pattern. Finally, the recovery period allowed 1 µM- exposed biofilms to reach the same levels of photosynthesis and biofilm formation potentials as control biofilms. Carbon fixation by prokaryotes was still more abundant for previously exposed biofilms (Figure 5).

**Figure 5:**
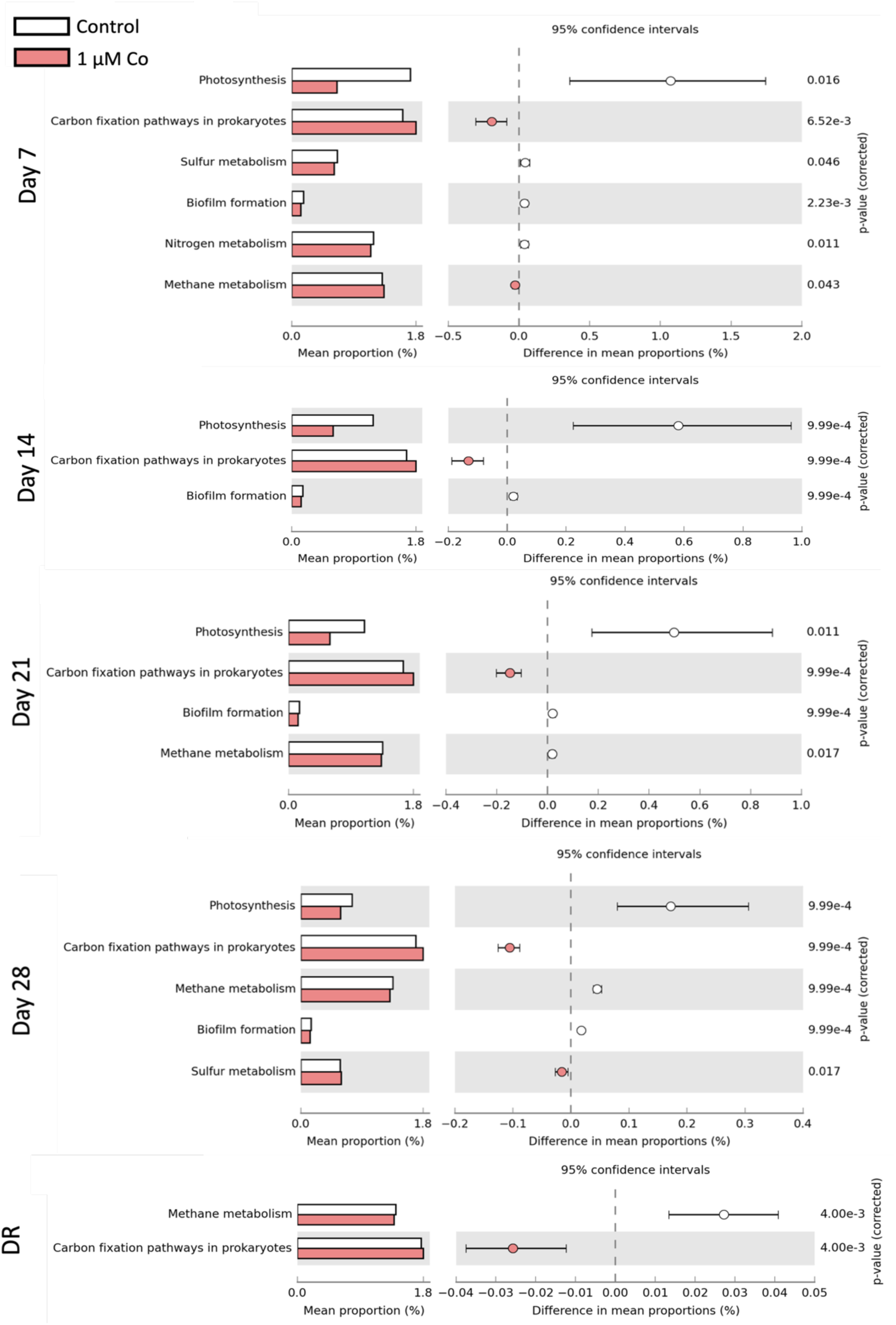
Predicted functional profile of prokaryotes within control biofilms and biofilms exposed to 1 µM of Co at each sampling time. Only the mean proportions and differences in mean proportions of pathways with significant difference of relative abundance (STAMPS comparison, White’s non-parametric test) are represented. DR: sampling day after 35 days of incubation without Co exposure.

## 5 Discussion

The present study aimed to assess the potential for prokaryotes within freshwater biofilms to be used as bioindicators for Co contamination in rivers. In a context where Co extraction and use has increased exceptionally in the last decades^1–3^, the health of aquatic ecosystems is under threat due to waste disposal in the environment or being impacted by nearby industrial sites^4,7^.

### 1. Co bioaccumulation in growing biofilms as a function of time and ambient concentrations

Cobalt concentrations within growing biofilms were well correlated with ambient free Co^2+^ concentrations, as recently observed in experiments with mature biofilms^21,22^ . Similar correlations were also found for metals such as Ni, Cd, Zn and Cu with biofilms, highlighting the potential for metal bioaccumulation in biofilms to be a good proxy for metal bioavailability^28,36–38^. While levels of total bioaccumulated Co were stable over biofilm formation, the internalized fraction gradually increased to represent, after 28 days of colonization, over 80% of the total bioaccumulated Co, regardless of the exposure concentration. It was in line with a recent study undertaken on mature biofilms where intracellular fraction of bioaccumulated Co increased over time, and represented the majority of bioaccumulated Co (71 ± 14%)^22^. After 28 days of exposure to 1 µM Co, comparable levels of intracellular Co were measure between studies (7.68 ± 1.10 and 8.81 ± 0.45 µmol.g_DW_^-1 22^) even if the Co bioaccumulation profile was more rapid and significant for mature biofilms. The higher bioaccumulation when exposed to higher Co was concomitant with higher prokaryotic abundances from D21 and a distinct microbial composition of biofilms, probably driven by selection of tolerant organisms.

### 2. Co impacts on the structure of bacterial community within growing biofilms

Biofilms developing in the *Gave de Pau River* are typical river biofilms dominated by *Proteobacteria, Bacteroidota, Planctomycetota* and *Cyanobacteria* phyla^15,18^. The formation of natural biofilms was characterized by a significant increase of proportions of *Planctomycetota* and a decrease of *Cyanobacteria*, probably linked to the decrease of light and temperature from October to December. Biofilms exposed to 0.1 µM Co showed similar prokaryotic community abundance, structure, and functional profile as control biofilms.

However, structural changes were observed in growing biofilms exposed to 0.5 µM and 1 µM Co. While the impact of metals on mature biofilms microbial communities has been widely described^22,38–41^, our study highlighted Co effects on early stages of biofilm colonization. Dissimilarities in the composition of the communities were already visible at D7, with this increasing with exposure time. These dissimilarities were due to shifts in the relative abundance of the major taxa as a function of Co concentrations. Among them, *Cyanobacteria* appeared extremely sensitive to Co for concentrations equal to or above 0.5 µM. *Cyanobacteria* sensitivity was also observed for environmental mature biofilms exposed to other metals (Ni, Cu, Zn and Pb)^20,42^. A limitation in the growth of an axenic *Cyanobacteria* strain for Co concentrations below 1 nM was also demonstrated^43^. This sensitivity highlights the potential for *Cyanobacteria* to be applied as a bioindicator for metal contamination. The high sensitivity of *Cyanobacteria* to Co likely led to the decrease of photosynthetic potential in the biofilms, with this already demonstrated elsewhere^44^. Such impacts may have repercussions on other heterotrophic bacteria since *Cyanobacteria* are important primary producers in biofilms^25^. Thus, the autotroph-heterotroph coupling, which is crucial for biofilm functioning and carbon cycling in aquatic ecosystems, could be impacted. However, the prediction of functional profile in the present study suggested that this crucial step of primary production for biofilms formation and maintenance was mainly supported by non-phototrophic CO_2_ fixators. This metabolic redundancy was in line with previous work which has described at least six alternatives to photosynthetic organisms for carbon fixation, including members of *Chlorobi*, *Chlorofexi*, *Aquaficea*, *Nitrospira*, *Planctomycetes, Proteobacteria* phyla^45–47^. Bacterial functional redundancy is well illustrated in the present study. Indeed, all biofilms reached similar dry biomass at maturity stage (D28), even though different prokaryotic community compositions were described.

*Planctomycetes* were also sensitive to Co, but to a lesser extent. Even if the relative abundance of this taxon increased over 21 days, its relative abundance was half that of the control biofilms. A recent study showed a decrease in the relative abundance of *Planctomycetes* in mature biofilms exposed to similar Co concentrations as tested here^22^. Abundances of *Planctomycetota* have been described to follow seasonal variations in river biofilms^48^, and a strong relationship between *Planctomycetes* class and macroalgal abundancies was observed^49^. Even if macroalgae were not investigated here, our data suggest that the lack of resources produced by photosynthetic primary producers in the presence of 0.5 and 1 µM Co induced a decrease of *Planctomycetota* members. Interestingly, some anaerobic ammonia-oxidizing members of *Planctomycetota* are able to perform the reductive acetyl-CoA pathway for carbon fixation, which might be one of the alternative pathways for the observed reduction of photosynthesis in the present study^45,50^. The acetyl-CoA pathway requires high levels of metals and coenzymes including cobalamin synthetized from Co^45^, suggesting that this pathway could be favored in biofilms exposed to high Co concentrations.

On the opposite, *Bacteroidota*, mainly composed of members of *Chitinophagales* and *Flavobacteriales,* were highly resistant and enriched in the presence of Co. *Bacteroidota* has high metabolic diversity and are able to degrade complex macromolecules. They are important degraders of suspended particles^15,16,51^. The increase of *Bacteroidetes* in the presence of 0.5 and 1 µM Co may be explained by their ability to degrade more refractory organic molecules within biofilms and ambient waters when labile organic compounds are less prevalent due to the sensitivity of *Cyanobacteria* sensitivity to Co.

### 3. Lack of resilience of prokaryotic communities after Co exposure

Our study also investigated the long-term effects of Co contamination after a return to the natural water composition. While prevention and detection are essential, it is also crucial to consider the resilience of exposed ecosystems to fully understand the extent of the impacts caused by metal contamination. The 35-day period after stopping Co injection did not allow a complete recovery of the biofilms when compared to the control biofilms. The levels of total bioaccumulated Co decreased but remained up to eight times higher in exposed biofilms than in control biofilms, with the prokaryotic communities also remaining different. The incomplete recovery of periphytic diatoms communities after a contamination of Cd and Zn^19^ or triclosan ^20^, has already been described. This lack of structural resilience in prokaryotic communities at the base of the trophic chain is likely to have vertical repercussions on the structure and interactions of higher levels^52^. Indeed, persistent imbalances on structure and key processes will impact resources and food availability such as oxygen and organic matter, affecting the food web and ecological status of rivers.

### 4. Prokaryotic communities are good bioindicators of a Co contamination

Even though microorganisms are key components of ecosystems, they are rarely used as biomonitoring tools. Microorganisms are known to have an important potential for bioindication since they are sensitive to environmental changes^39,53,54^. Moreover, their activity and position as the basis of the trophic chain allow them to act as an early warning tool before repercussions at higher trophic levels are observed. The present study aimed at evaluating the potential of prokaryotic communities to be used as a bioindicator tool of Co exposure. The sensitivity of major taxa, such as *Cyanobacteria* and *Planctomycetes* or the resistance of others, such as *Bacteroidetes,* were well reflected with the microorganisms rapid response to disturbance in rivers. In addition, targeting prokaryotic communities with a molecular approach allows for the potential prediction of the functional repercussions the exposure to a contaminant can cause. They can also be useful in a context of evaluating the restoration of aquatic ecosystems as the history of Co contamination was still visible on the prokaryotic community structure after the conclusion of the stress induction and a following recovery period. Prokaryotic communities studied in the present work demonstrated, thus, evidence as viables alternatives to traditional methods of counting and morphological analyses, whose accuracy is often affected by human biases^55^. Indeed, microbial approaches are less time-consuming and labor-intensive if used as a routine method. Another advantage is that they offer a complete picture of the impacts of a river contamination from communities’ structure and groups interactions to functional profile, which also provides deeper insights into the repercussions which may occur at higher trophic levels and even at the ecosystems level. In 2009, bacteria and benthic invertebrates bioindication ability were compared and bacteria allowed to detect only the most impacted sites by comparison with macroinvertebrates^56^. The authors suggested that more research is required before endorsing the use of bacterial community as an indicator of freshwater ecological health. However, since 2009, major technological advances in molecular techniques have been achieved, allowing to a deeper analysis of the microbial communities. There is an enormous potential for their use as a monitoring tool, alone or by their integration with other physical, chemical and biological measures in a modelling approach, to develop a better understanding of metal impacts on environmental health^57^. As with most bioindicator methods, the most challenging part of the process is the availability of local natural biofilm composition reference data of since water composition can vary depending on seasons, river conditions and geography, and the influence these factors have on the natural structure of reference biofilms. Further studies focusing on sensitive and resistant groups described in the present work are needed to confirm their behavior in the presence of metals. Also, the interactions between all compartments within biofilms including microalgae should be considered in the future to assess metal impact on the whole riverine ecosystem.

## Acknowledgment

We thank the members of TotalEnergies Environment & Sustainable Development Team at Pôle d’Études et de Recherches (PERL, Lacq, France) for the access to pilot rivers facilities and experimental assistance during the mesocosm experiment. We also thank Claire Gassie (Université de Pau et des Pays de l’Adour, E2S-UPPA, CNRS, IPREM, Pau, France) for her technical assistance during laboratory work and characterization of microbial communities. This research was funded by the Partnership Chair E2S-UPPA-TotalEnergies-Rio Tinto (ANR- 16-IDEX-0002).

## Notes

### Competing Interest Statement

The authors have declared no competing interest.

