## Supplementary Materials and Figures for "Cobalt effects on prokaryotic communities living in growing river biofilms: impact on their colonization kinetics, structure and functions"

Includes:

-Supplementary material and methods

-7 supplementary figures

### Supplementary material and methods

### 1.Water analysis

Before biofilm collection, the main physico-chemical parameters of water were measured in each stream: pH, temperature, dissolved oxygen, and conductivity using a multi-parametric probe (HACH, IA, USA). Water was filtered on 0.45 µm polysulfone encapsulated filter membrane (Minisart®, Supelco, Germany) and collected in polypropylene tubes (Metafree®, Labcon, North America) for dissolved Co, cation, and anion analyses. Cobalt concentrations in exposure media were analyzed by Inductively Coupled Plasma Mass Spectrometry (ICP-MS model 7500/7700, Agilent, CA, USA). Anion concentrations were measured by ion chromatography (Dionex Aquion ®, Thermo-Fisher, MA, USA) and cations by ICP-AES (iCAP 6500 Thermo-Fisher, MA, USA). SLR-6 was used as a certified material to assess the accuracy of ICP-MS (recovery = 103 ± 5% for Co, n = 5) and ICP-AES analysis (recovery of 98 ± 1% for Ca^2+^, n = 3). For DOC measurements, an additional 125 mL of water was filtered water on microfiber 0.7 µm GF/F filters (Whatman Cytiva, USA), and collected in amber borosilicate containers previously burnt at 450°C. DOC concentrations were determined by Shimatzu TOC-L device (Japan). All the analyses were performed in triplicates.

**2.Major biofilm characteristics (fresh and dry weight and chlorophyll content)**

For chlorophyll content analysis, biofilms were freeze-dried for 24 hours and the pigments were extracted in the dark using 10 mL of 90% acetone. After 40 minutes of sonication, samples were incubated in the dark overnight at 4°C and then centrifugated at 6,000 rpm for 10 minutes. The chlorophyll contents were measured in the supernatant by spectrophotometry with an absorbance measure from 700 to 400 nm (Lambda 750, PerkinElmer, MA, USA). Concentrations of chlorophyll *a*, *b* and *c1+c2* were then calculated^2^.

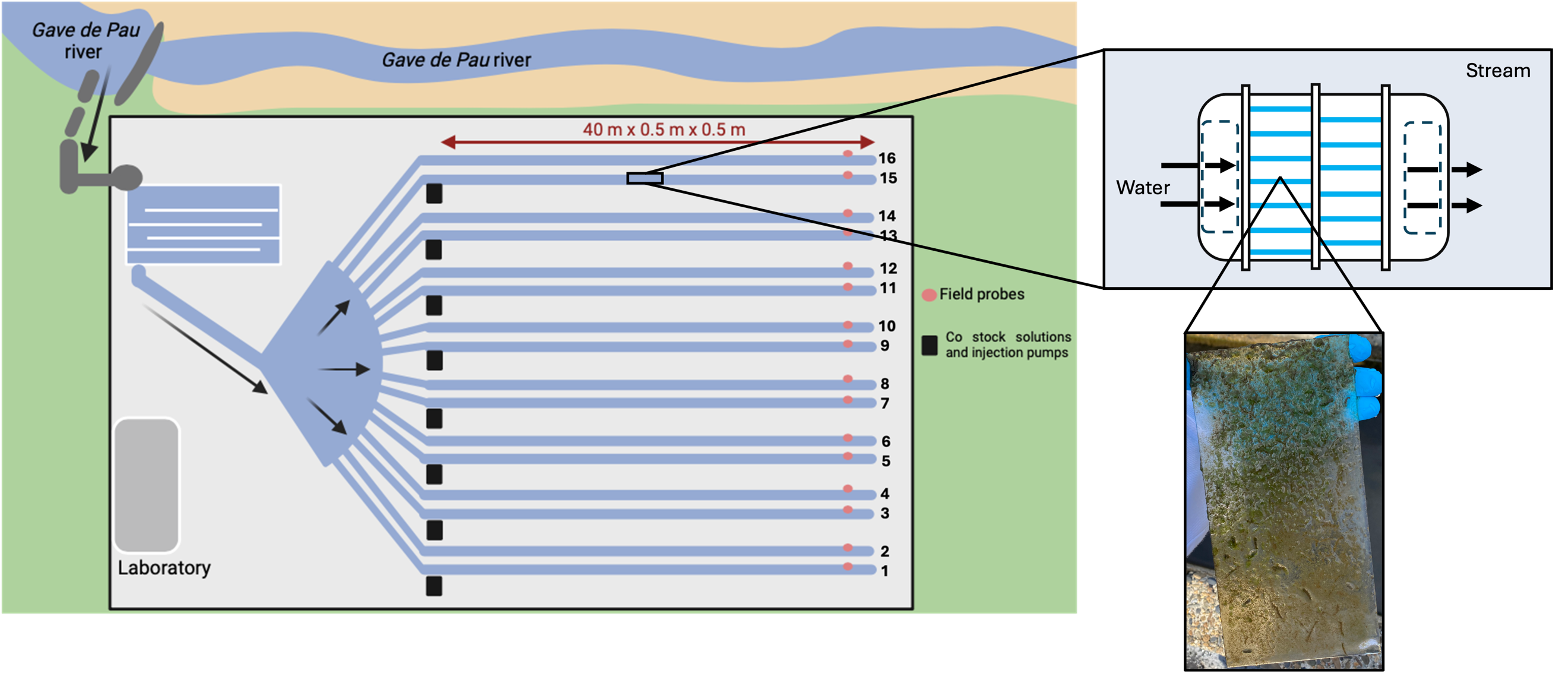

**Figure S1**: The pilot rivers facility is an outdoor mesocosm located in Lacq, France and hosted by TotalEnergies^1^. The facility consists of 16 artificial streams (40 m long, 0.5 m wide, 0.5 m deep and with a flow rate of 7.5 m^3^.h^-1^) connected to the *Gave de Pau River* as an open circuit. For the experiment, twelve channels were selected, and cobalt concentrations were randomly assigned as follows: three replicate channels for each exposure conditions (0.1 µM Co: streams 5, 10 and 15; 0.5 µM Co: streams 2, 3 and 9; 1 µM Co: streams 7, 11 and 12). Solutions of cobalt (Cobalt(II) chloride hexahydrate, 98%, Thermo Scientific Chemicals) were injected continuously (71 mL.h^-1^) in channels with piston pumps (Netzch, Nemo, Germany), starting one day before the experiment and for a total period of 29 days. *Gave de Pau* water was used in control stream (6, 8 and 14) in triplicates with background concentrations of cobalt. Field probes (Hach, IA, USA) located at each end of channel were used to measure Temperature (°C), pH, conductivity, oxygen saturation and dissolved oxygen before sampling. A box with 13 glass slides was placed in each stream for the experiment at a distance of 15 meters from the injection pumps, as support for biofilms colonization.

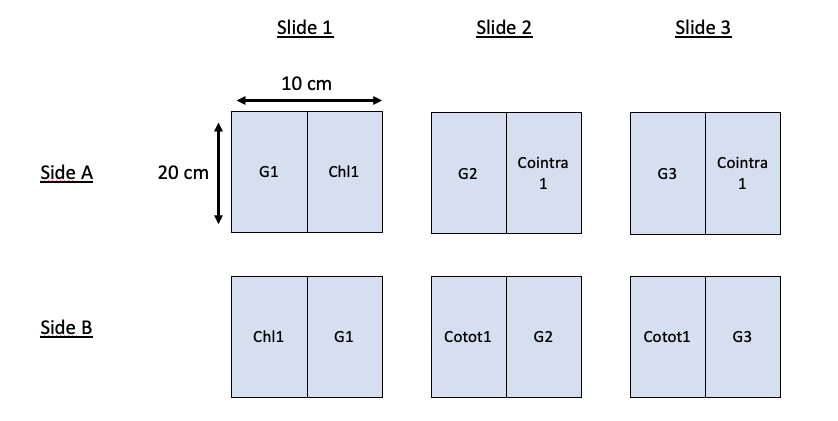

**Day 7 and Day 14**

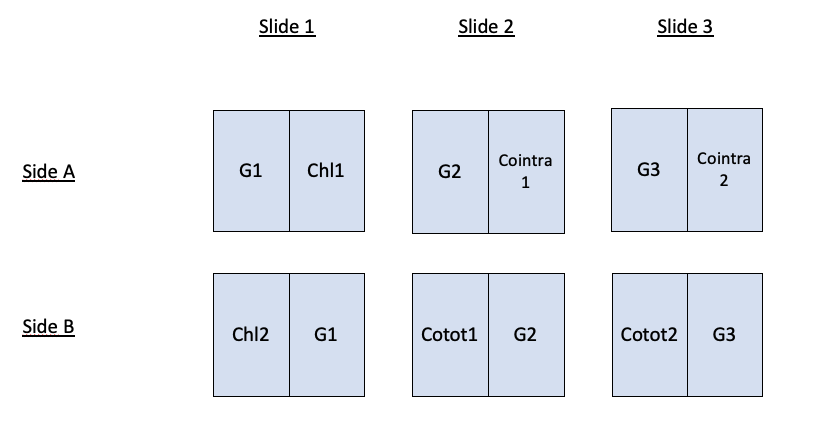

**Day 21 and Day 28**

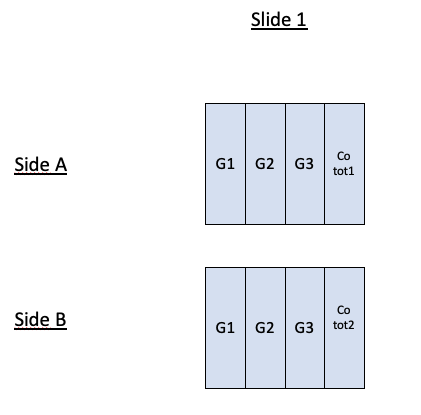

**After *recovery***

**Figure S2:** Detailed plan of the glass slides used for biofilm collection throughout the mesocosm experiment. Thirteen sterile blank glass slides (10 x 20 cm) were submerged in each channel. At each sampling time (from D7 to D28) and for each channel, three slides (6 sides) were used for biofilm collection. For biofilm sampling after the recovery period, one glass slide (two sides) was used for biofilm collection. Glass slides were scratched with sterile microscope slides. Samples were named according to the analyses they were dedicated to: chlorophyll content (Chl), DNA extraction (metagenomic, G), total (Cotot) and intracellular (Cointra) cobalt bioaccumulation and the number indicates the replicate (1-3). Three replicates by streams were sampled during cobalt exposure for metagenomic analysis. At D7 and D14, one replicate per stream was collected for chlorophyll and cobalt bioaccumulation (one for intracellular and one for total) analyses because of the low quantity of biofilm. Two replicates were then collected at D21 and D28 for chlorophyll and cobalt bioaccumulation analyses. After DR, three replicates were also collected for metagenomic analysis and remaining biofilms on slides were collected to obtain two replicates for total cobalt bioaccumulation analyses.

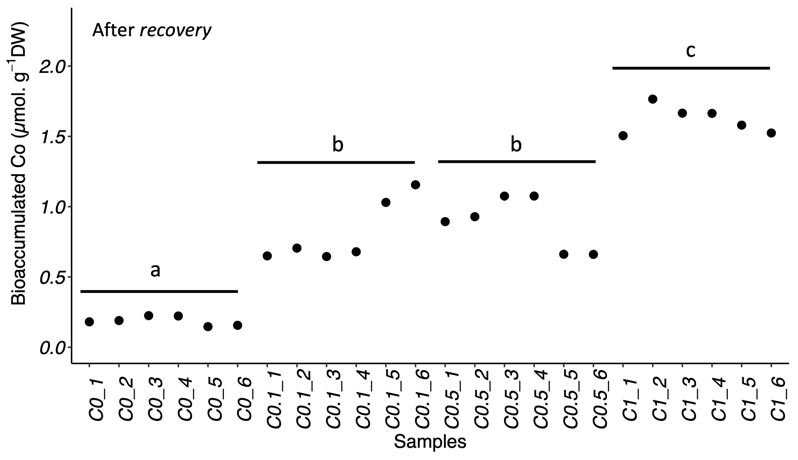

(C)

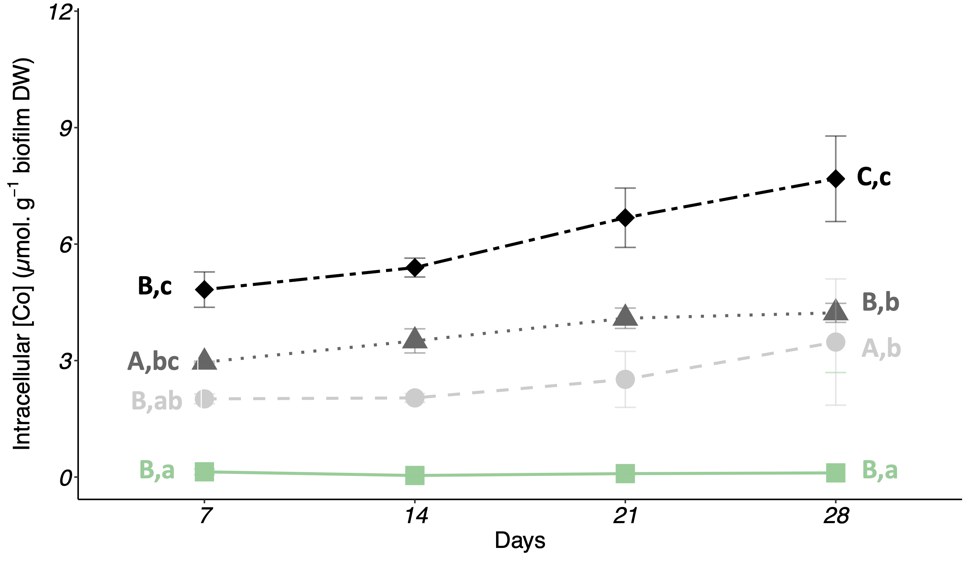

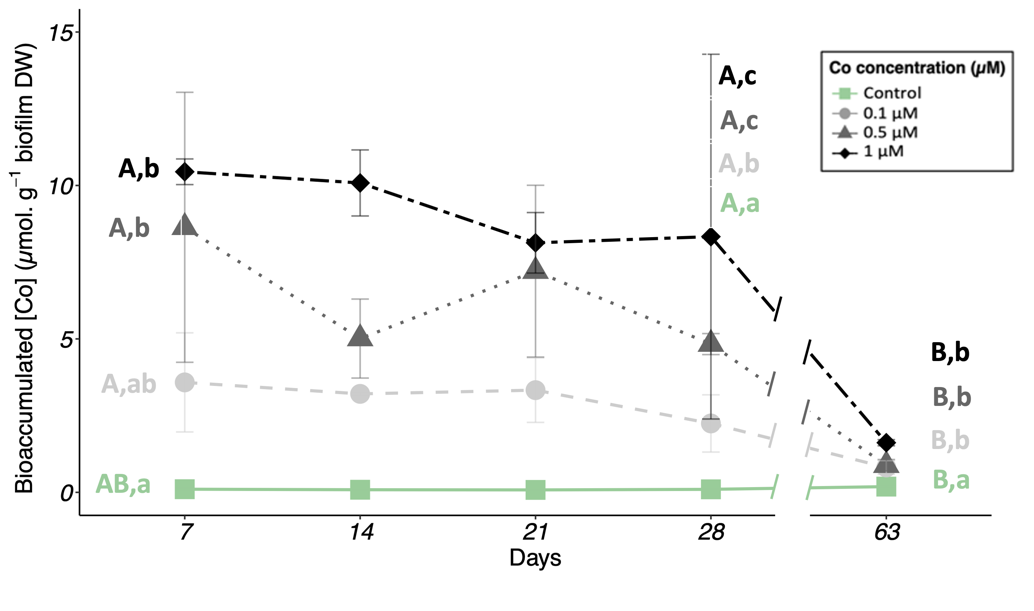

(B)

(A)

**Figure S3:** Kinetics of (A) total and (B) internalized Co accumulation in biofilms as a function of time for each exposure condition. Colors indicate the increasing Co concentrations by comparison with control conditions (~~■~~: Control conditions; ~~•~~: 0.1 µM Co; ~~▲~~: 0.5 µM Co; ~~♦~~: 1 µM Co). (C) Levels of total bioaccumulated cobalt after 35 days of recovery (DR). Samples are names according to the exposure condition of biofilms and the number of replicates: Control (C0); 0.1 µM Co (C0.1); 0.5 µM Co (C0.5); 1 µM Co (C1) and replicates from 1 to 6 for each exposure condition. Bioaccumulation data include samples in one replicate within a stream for Day 7 and Day 14, and two replicates within a stream for Day 21, Day 28 and DR.

a, b, c, d Significant groups defined from post-hoc Dunn’s test after Kruskall-Wallis non-parametric test between Co concentrations at a sampling time.

A, B, C, D Significant groups defined from post-hoc Dunn’s test after Kruskall-Wallis non-parametric test between sampling times for a concentration.

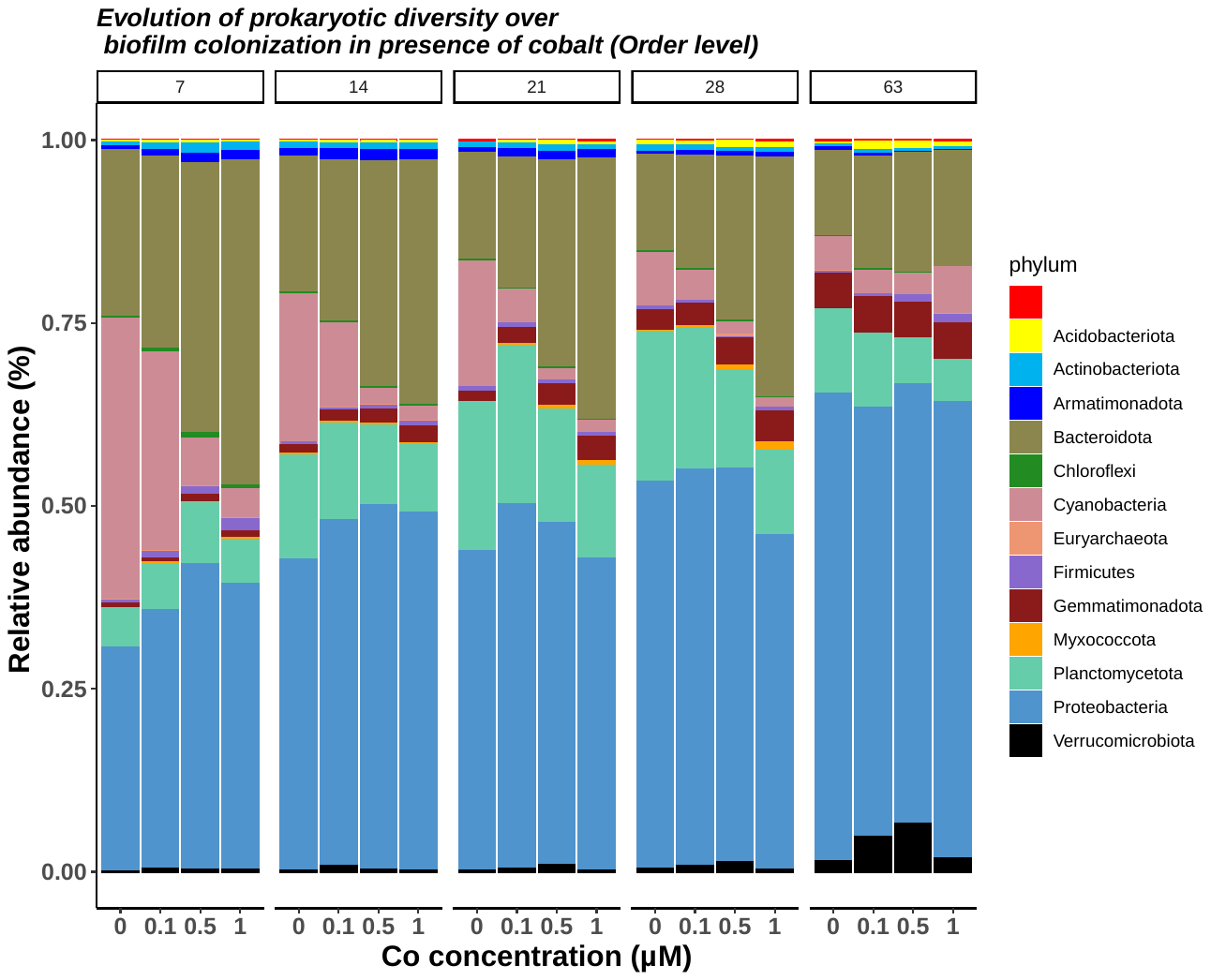

(A)

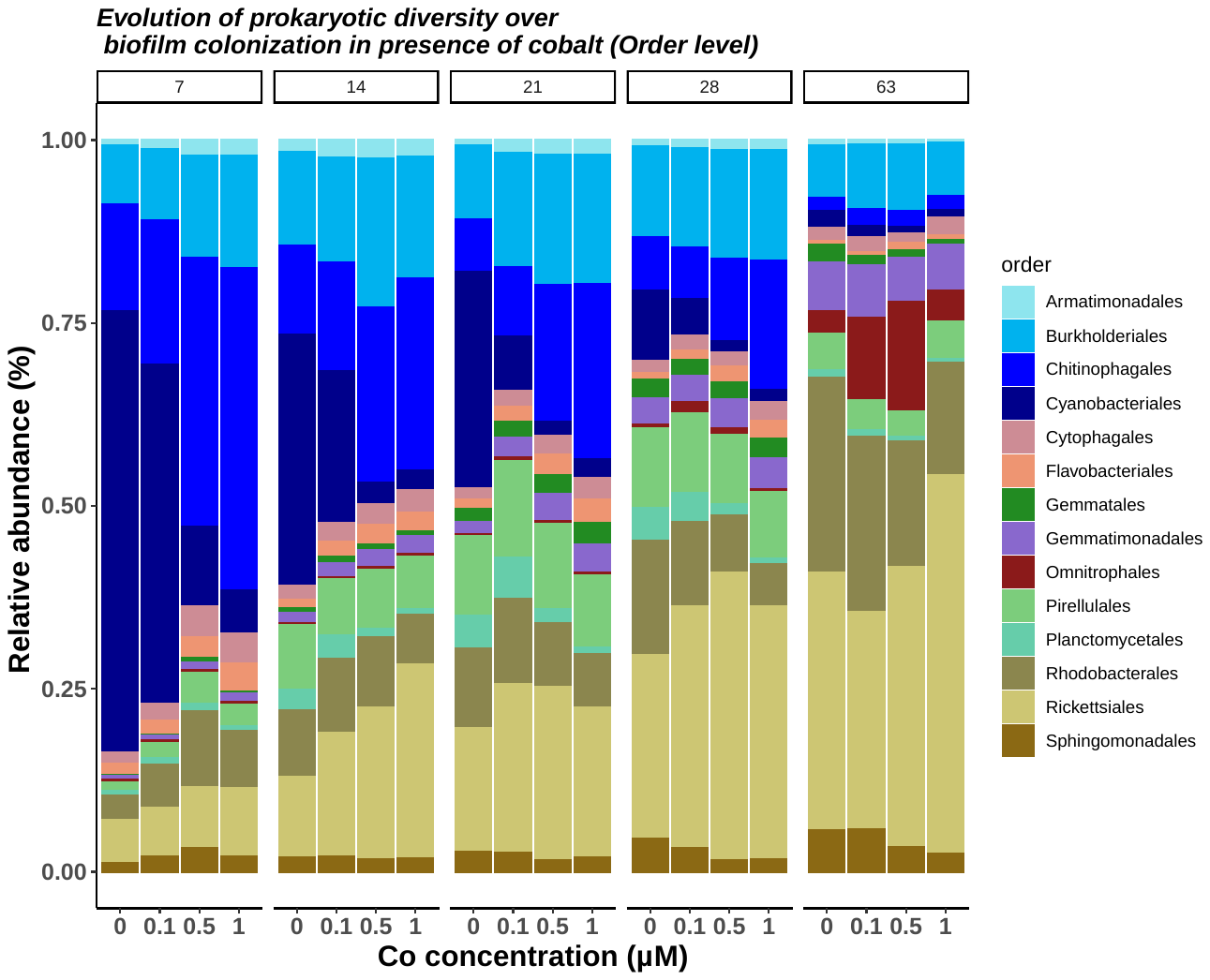

(B)

**Figure S4**: Prokaryotic community composition at phylum level (A) and family level (B) as a function of time and Co concentration.

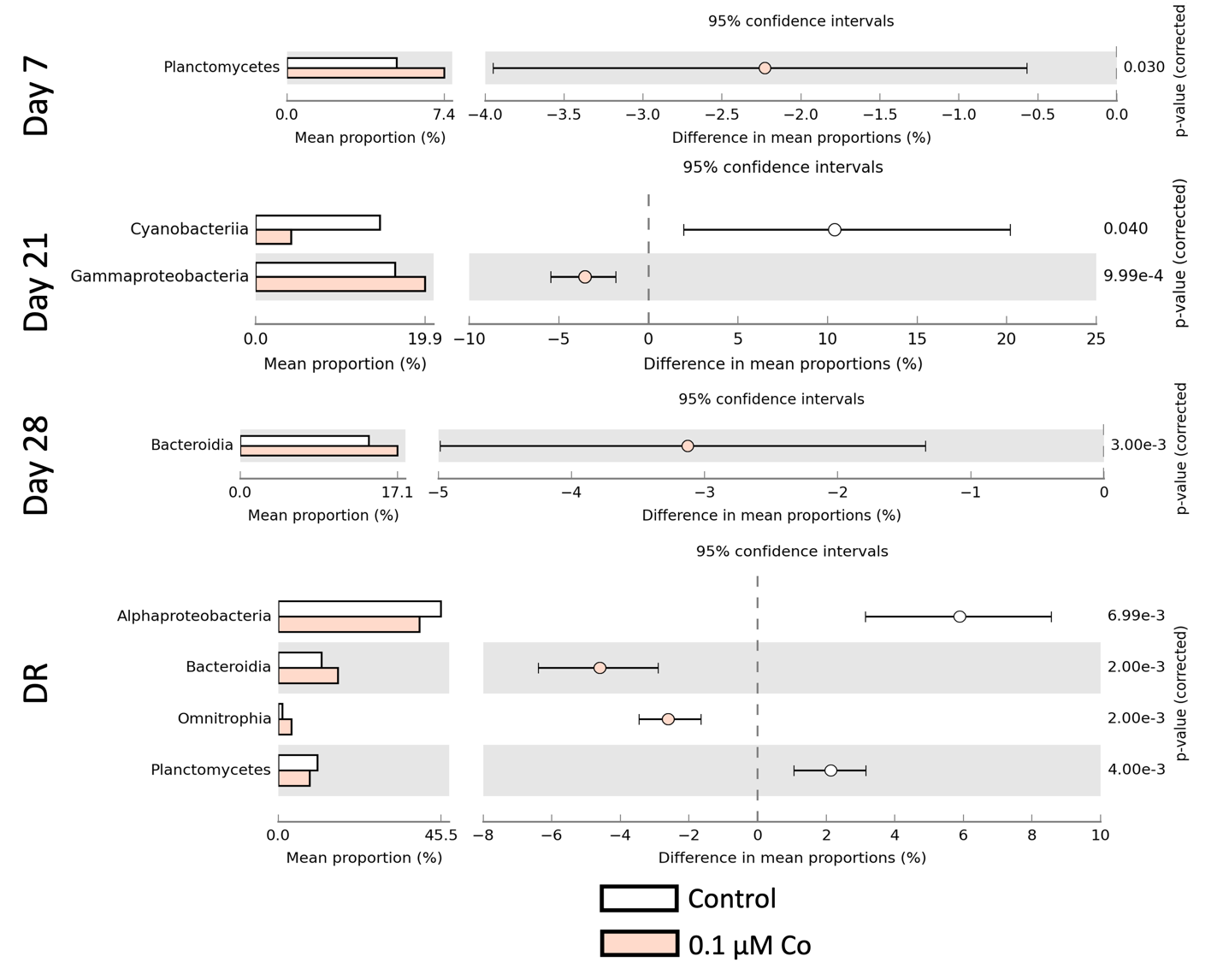

**Figure S5:** Mean proportions and differences in the mean proportions of taxa with statistically significant differences (White’s non-parametric test) between control and 0.1 µM-exposed biofilms based on STAMPS comparison. In white, control condition, in red, biofilms exposed to 0.1µM of Co.

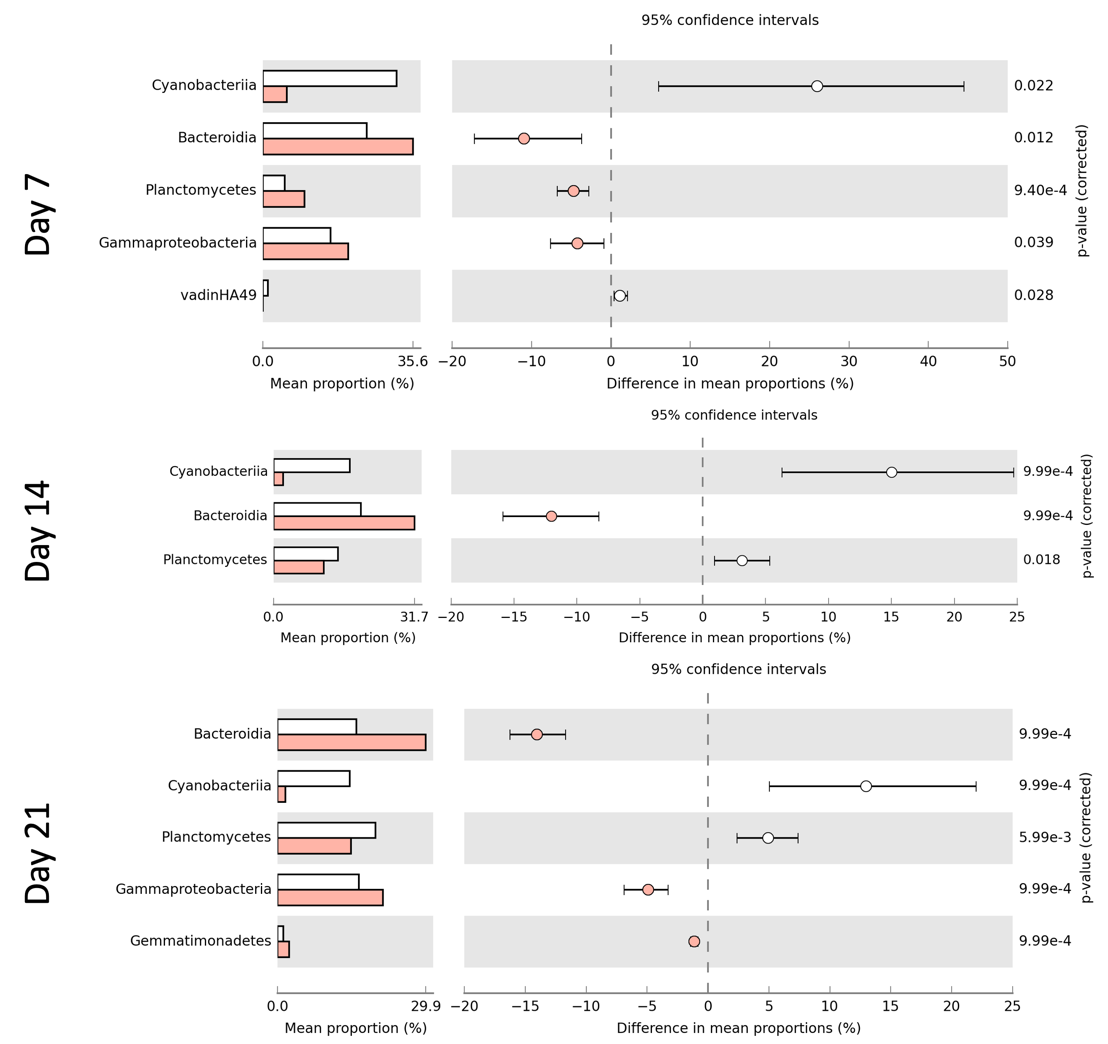

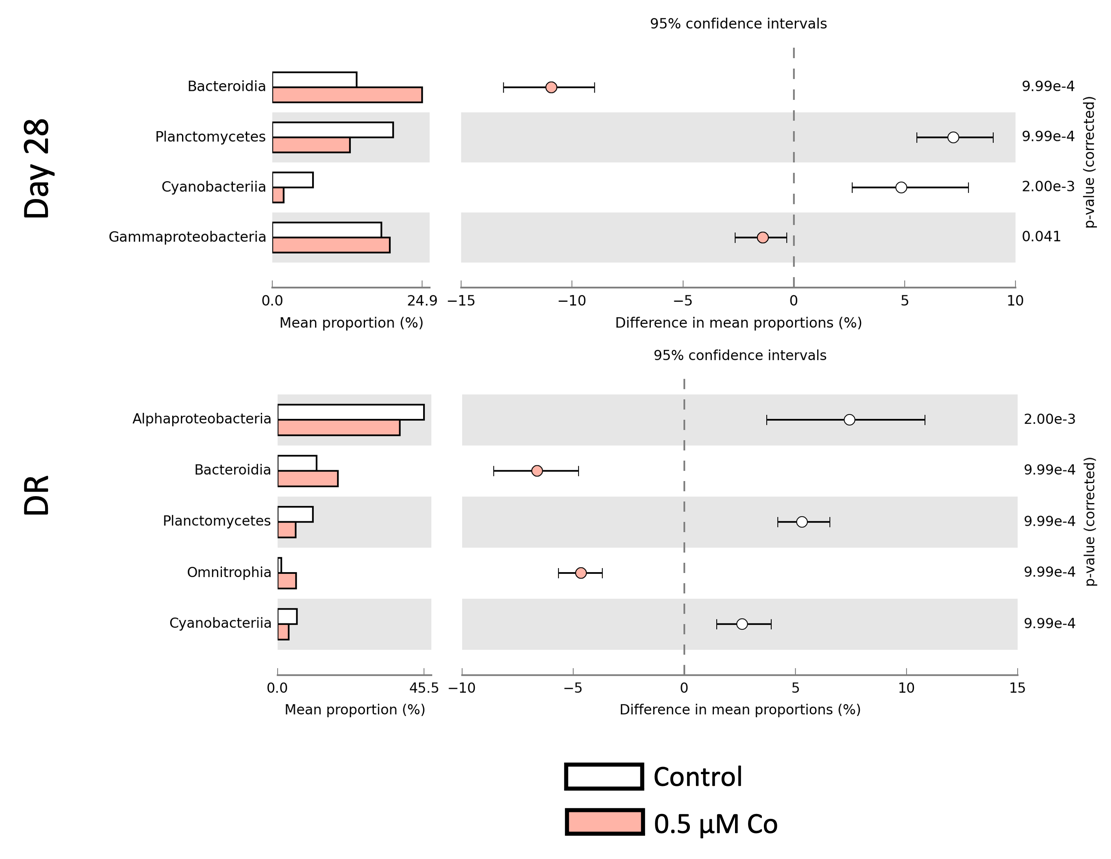

**Figure S6**: Mean proportions and differences in the mean proportions of taxa with statistically significant differences (White’s non-parametric test) between control and 0.5 µM-exposed biofilms based on STAMPS comparison. In white, control condition, in red, biofilms exposed to 0.5µM of Co.

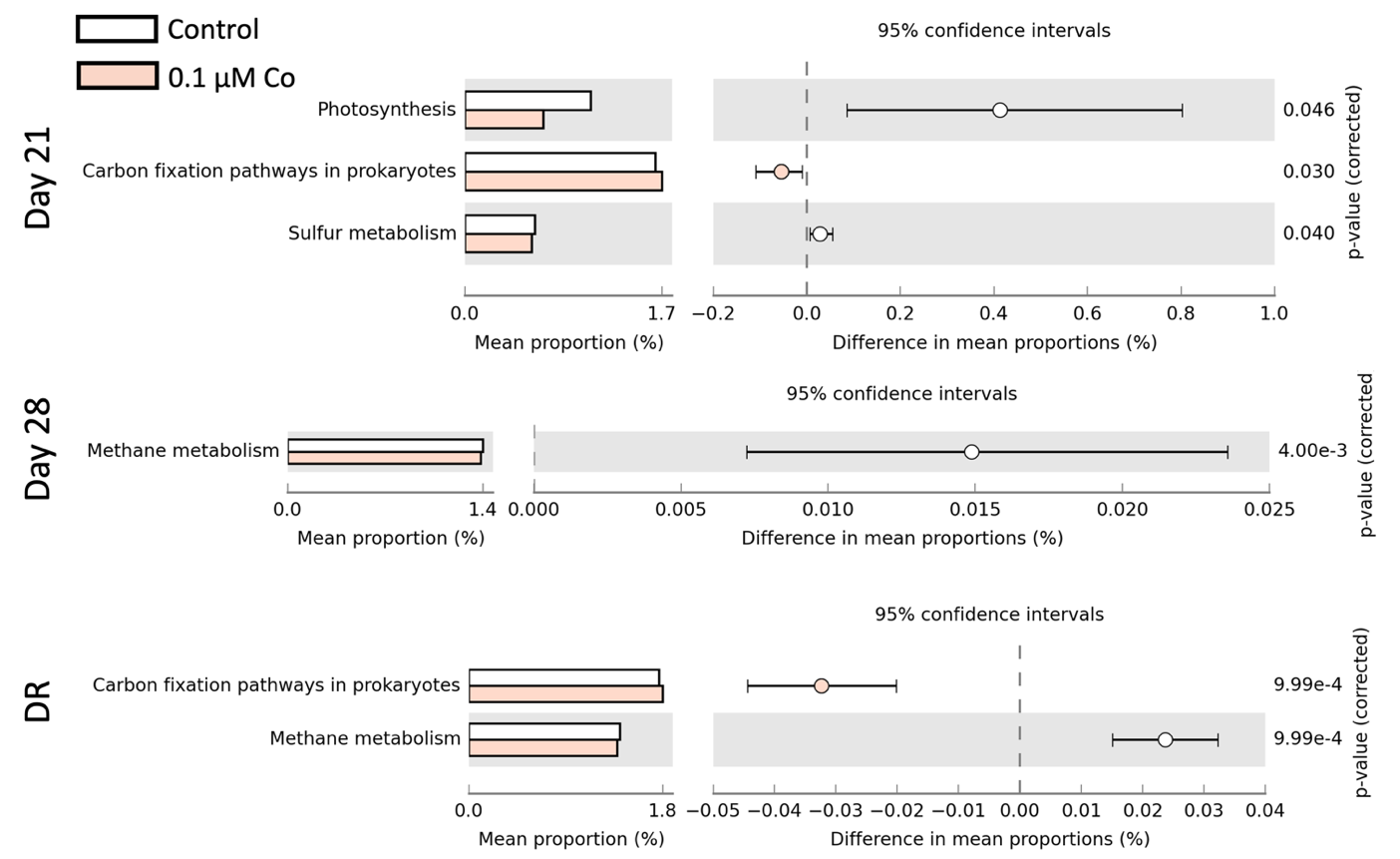

**Figure S7:** Predicted functional profile of prokaryotes within control biofilms and biofilms exposed to 0.1 µM of Co at each sampling time. Only the mean proportions and differences in mean proportions of pathways with significant difference of relative abundance (STAMPS comparison, White’s non-parametric test) are represented. DR: sampling day after 35 days of recovery without Co exposure. In white, control condition, in red, biofilms exposed to 0.1µM of Co.

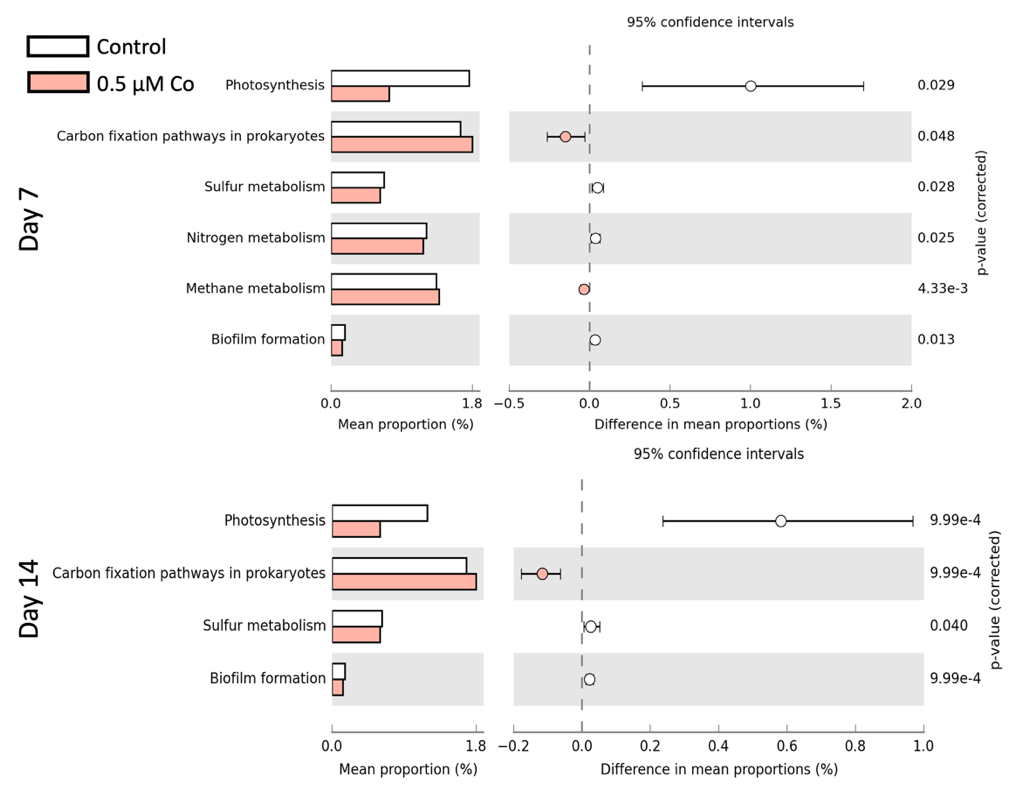

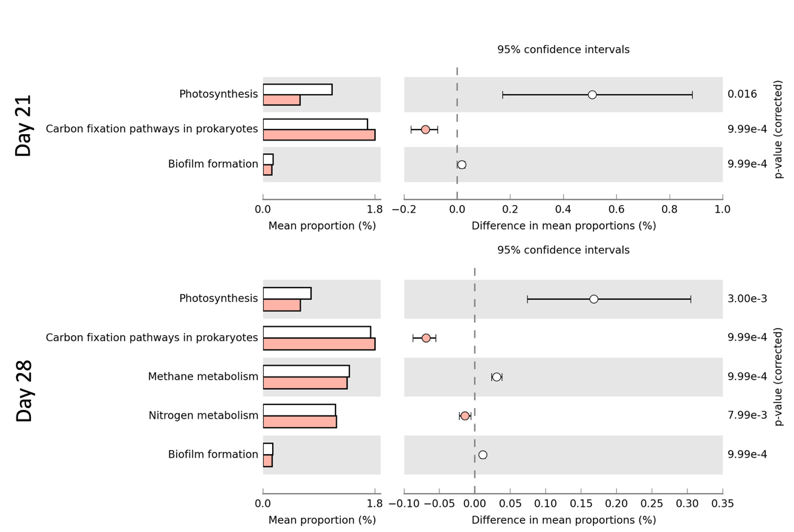

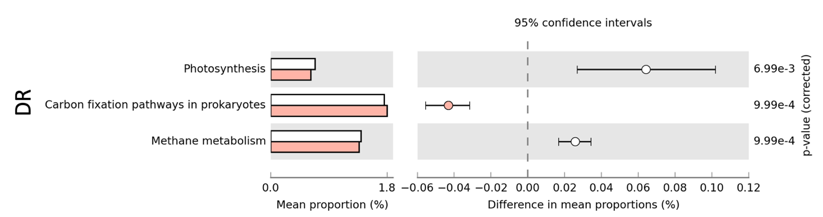

**Figure S8:** Predicted functional profile of prokaryotes within control biofilms and biofilms exposed to 0.5 µM of Co at each sampling time. Only the mean proportions and differences in mean proportions of pathways with significant difference of relative abundance (STAMPS comparison, White’s non-parametric test) are represented. DR: sampling day after 35 days of recovery without Co exposure. In white, control condition, in red, biofilms exposed to 0.1µM of Co.

**Supplementary tables:**

**Table S1:** Physicochemical parameters measured in the pilot rivers waters at each sampling time.

**Table S2:** Total and internalized concentrations of Co bioaccumulated by growing biofilms for 28 days and after the recovery period (Total only). A, B, C, D Significant groups defined from post-hoc Dunn’s test after Kruskall-Wallis non-parametric test between sampling times for a concentration.

a, b, c, d Significant groups defined from post-hoc Dunn’s test after Kruskall-Wallis non-parametric test between Co concentrations at a sampling time.

**Table S3:** Biological parameters of collected biofilms throughout the colonization period for each condition of exposure.

**Table S4:** Indexes of alpha diversity of prokaryotic community within biofilms colonized in presence of cobalt.

**Table S5:** Cobalt effect on beta-diversity of prokaryotic communities over biofilms colonization in presence of Co. Group comparison scores and significances levels defined after Pairwise permanova test. Sampling times and Co concentration of exposure are annotated as follows: T1 (Day 7), T2 (Day14), T3 (Day21), T4(Day28), T5(Recovery), C1 (Control), C2(0.1µM Co), C3 (0,5 µM Co), C4 (1 µM Co)

Supplementary materials references:

(1)	Cailleaud, K.; Bassères, A.; Gelber, C.; Postma, J. F.; Ter Schure, A. T. M.; Leonards, P. E. G.; Redman, A. D.; Whale, G. F.; Spence, M. J.; Hjort, M. Investigating Predictive Tools for Refinery Effluent Hazard Assessment Using Stream Mesocosms. *Enviro Toxic and Chemistry* **2019**, *38* (3), 650–659. https://doi.org/10.1002/etc.4338.

(2)	Jeffrey, S. W.; Humphrey, G. F. New Spectrophotometric Equations for Determining Chlorophylls a, b, C1 and C2 in Higher Plants, Algae and Natural Phytoplankton. *Biochemie und Physiologie der Pflanzen* **1975**, *167* (2), 191–194. https://doi.org/10.1016/S0015-3796(17)30778-3.
